# Genetic drift vs natural selection affecting the evolution of spectral and functional traits of two key macrophytes: *Phragmites australis* and *Nuphar lutea*

**DOI:** 10.1101/2023.06.19.545543

**Authors:** Maria Beatrice Castellani, Alice Dalla Vecchia, Rossano Bolpagni, Roberto Natale, Erika Piaser, Lorenzo Lastrucci, Andrea Coppi, Paolo Villa

**Affiliations:** Department of Biology, University of Florence, via Micheli 1, 50121 Florence, Italy; Department of Chemistry, Life Sciences and Environmental Sustainability, University of Parma, Parma, Italy; Institute for Electromagnetic Sensing of the Environment (IREA), National Research Council of Italy (CNR), Via A. Corti 12, 20133 Milano; Natural History Museum, Botanical Collections, University of Florence, via G. La Pira 4, 50121 Firenze, Italy

**Author notes:** Andrea Coppi and Paolo Villa should be considered joint senior authors.

**Keywords:** aquatic plants, freshwater ecosystems, common reeds, yellow water-lily, *P*_st_ - *F*_st_ comparison, leaf reflectance

## Abstract

Both genetic and phenotypic intraspecific diversity play a crucial role in the ecological and evolutionary dynamics of organisms. Several studies have compared phenotypic divergence (*P*_st_) and differentiation of neutral loci (*F*_st_) to infer the relative roles of genetic drift and natural selection in population differentiation (*P*_st_ - *F*_st_ comparison). For the first time, we assessed and compared the genetic variation and differentiation at the leaf trait level in two key macrophytes, *Phragmites australis* and *Nuphar lutea*.

To this aim, we quantified and described the genetic structure and phenotypic diversity of both species in five lake systems in north-central Italy. We then investigated the relative roles of genetic drift and natural selection on leaf trait differentiation (*P*_st_ - *F*_st_), assuming that *F*_st_ reflects divergence caused only by genetic drift while *P*_st_ also incorporates the effects of selective dynamics on the phenotype.

In terms of genetic structure, the results for *P. australis* were in line with those observed for other Italian and European conspecific populations. Conversely, *N. lutea* showed a more complex genetic structure than expected at the site level, likely due to the combined effect of genetic isolation and its mixed mating system. Both species exhibited high variability in leaf functional traits within and among sites, highlighting a high degree of phenotypic plasticity. *P*_st_ - *F*_st_ comparisons showed a general tendency towards directional selection in *P. australis* and a more complex pattern in *N. lutea*. Indeed, the drivers of phenotypic differentiation in *N. lutea* showed a variable mix of stabilizing and directional selection or neutral divergence at most sites.

The prevalence of vegetative over generative reproduction leads *P. australis* populations to be dominated by a few clones that are well adapted to local conditions, including phenotypes that respond plastically to the environment. In contrast, in *N. lutea* the interaction of a mixed mating system and geographical isolation among distant sites tends to reduce the effect of outbreeding depression and provides the genetic basis for adaptive capacity.

The first joint analysis of the genetic structure of these two key macrophytes allowed a better understanding of the relative roles of genetic drift and natural selection in the diversification of phenotypic traits within habitats dominated by *P. australis* and *N. lutea*.

## 1. Introduction

In the last decades, several studies on evolutionary processes inferred the relative role of genetic drift and natural selection on species diversification (Orsini *et al*., 2013; Andrews *et al*., 2016). However, the extent to which both evolutionary forces affect populations depends largely on the ecological features of species. It is well known that the degree of individual specialization varies widely among plant species, mirroring a multitude of physiological, behavioral, and ecological mechanisms (Bolnick *et al*., 2003) directly involved in driving the species’ response to both biotic and abiotic factors (Hughes *et al*., 2008; Bolnick *et al*., 2011; Eller *et al*., 2017) and their ecological resilience (Moran *et al*., 2015). Therefore, a better understanding of patterns and processes of genetic and phenotypic diversity at intraspecific level is crucial for ecological, evolutionary and conservation studies (Chave, 2013; Mimura *et al*., 2017).

To determine how the degree of genotypic and phenotypic differentiation is caused by selective versus neutral processes, several studies – as far as we know, none strictly on aquatic plants – have compared phenotypic divergence (*P*_st_) and differentiation of neutral loci (*F*_st_), i.e. *P*_st_ – *F*_st_ comparisons (Brommer, 2011; Leinonen *et al*., 2013). If phenotypic features evolved neutrally, the proportion of their variation among populations should be comparable to that of variation in allele frequencies at neutral loci (*P*_st_ = *F*_st_). On the other hand, if *P*_st_ is higher or lower than *F*_st_, the differentiation of phenotypic features is more likely shaped by natural selection (directional or divergent), or stabilizing selection, respectively (Merilä and Crnokrak, 2001; Leinonen *et al*., 2008; Whitlock, 2008; Chapuis *et al*., 2008; Martin *et al*., 2008; Brommer, 2011; Leinonen *et al*., 2013; Seymour *et al*., 2019; Marin *et al*., 2020). According to Leinonen *et al*. (2006) and Brommer (2011), *P*_st_ is a proxy of the quantitative genetic differentiation index (*Q*_st_) and it is used when additive variance cannot be readily easily quantified (i.e., in field studies; Brommer, 2011). The problems of using *P*_st_ as an approximation of *Q*_st_ are well known in the literature. However, the estimation of the *Q*_st_ index requires individuals from different populations to be grown together in a common garden (Leinonen *et al*., 2008), so the *P*_st_ index is commonly used when the populations under study are located in different areas and may present locally adapted forms.

Regarding the interconnection between genetic differentiation and phenotypic plasticity, a quantitative focus can be provided by measurement of variation in plant functional traits (FTs), especially in relation to those pertaining to the leaf economics spectrum (Pierce *et al*., 2012; Díaz *et al*., 2016; Dalle Fratte *et al*., 2019a; Dalla Vecchia *et al*., 2020). FTs such as leaf total area (LA, including petioles), specific leaf area (SLA) or pigments content (chlorophyll-a, Chl-a; and the ratio between chlorophyll-a and -b, Chl-a/Chl-b) help to quantify the responses of species and communities to abiotic factors. Moreover, new windows on plant functional ecology were opened by the expansion of remote sensing-based applications, enabling high-throughput investigation of plant variability in spatial and temporal dimensions (Wang and Gamon, 2019), recently extended to account for specific spectral features of aquatic plant species (Villa *et al*., 2021). Joining functional and spectral-based approaches for characterizing plant structure and physiology across sites and ecosystems can offer a cross-feedback for a better understanding of ecosystem functioning and consequently providing effective management strategies (Villa *et al*., 2017, Castellani *et al*., 2023).

Here, we studied two key freshwater macrophytes, the helophyte *Phragmites australis* (Cav.) Trin. Ex Steud. (or common reed) and the floating hydrophyte *Nuphar lutea* (L.) Sm. (or yellow water-lily), as models to infer the relative role of genetic drift and natural selection on the diversification of leaf traits. Both target species are often dominant and widespread macrophytes and as such can act as ecosystem engineers, shaping the colonized habitats through their physiology and physical structure (Thomaz, 2021). Common reed is a sub-cosmopolitan helophyte, dominating the riparian vegetation of most freshwaters and brackish wetlands globally. This species may also colonize disturbed wetlands, artificial ditches, mined areas, and landfill, thus proving a broad ecological amplitude. These characteristics make *P. australis* a keystone species in freshwater ecosystems and support several habitats and non-habitat ecosystem services (Kiviat, 2013). *P. australis* plays an essential role as a wind and wave breaker (Takeda and Kurihara 1988; Vymazal, 2011; Karstens *et al*., 2016) and its functional traits seem to be determined by trophic conditions (Eid *et al*., 2021). Yellow water-lily is a floating-leaved macrophyte distributed across lower latitudes of Europe, northwest Africa (locally known for Algeria), and eastwards to central and southwest Asia (Padgett, 2007). Its robust petiole and the large leaf area foster a low hydrodynamic environment mitigating the wave movement and promoting sedimentation (Puijalon *et al*., 2011; Schoelynck *et al*., 2014). Its leaf traits appear to be regulated by water depth and sediment features in hyper-eutrophic environments (Dalla Vecchia and Bolpagni, 2022).

The main aim of this work was to assess and compare the genetic variation and leaf trait differentiation of the two target macrophytes, *P. australis* and *N. lutea*. We first quantified and described the genetic structure and phenotypic diversity – estimated from leaf functional traits directly measured and inferred from foliar reflectance – in different populations sampled across lake systems in northern and central Italy. We then investigated the correspondence between the degree of population differentiation at neutral genetic markers and quantitative FTs, respectively expressed by *F*_st_ and *P*_st_ indices, to infer the forces (i.e., stabilizing selection, directional/divergent selection, or neutral divergence) shaping the population differentiation of each trait. Comparing *F*_st_ and *P*_st_, we assume that *F*_st_, determined by neutral markers, reflects divergence caused only by genetic drift (Reynolds *et al*., 1983). Thus, *F*_st_ provides a null expectation and allows estimation of what would happen to populations in the absence of selection (Merilä and Crnokrak, 2001; Noguerales *et al*., 2016; but see Edelaar *et al*., 2011). On the other hand, *P*_st_ also incorporates the effects of selective dynamics on the phenotype. Consequently, the *P*_st_ - *F*_st_ comparison involves three possibilities in deciphering the evolution of sampled populations of *P. australis* and *N. lutea*: 1) *P*_st_ > *F*_st_ indicates that traits divergence overcome that attributable to genetic drift alone, suggesting that directional or divergent selection (DS) is leading to population differentiation; 2) *P*_st_ = *F*_st_, indicates that neutral divergence cannot be excluded as a possible cause of phenotypic differentiation (NS); 3) *P*_st_ < *F*_st_ indicates that phenotypic differentiation is less than expected based on genetic drift alone, suggesting that trait divergence is most likely the product of stabilizing selection (SS) (Merilä and Crnokrak, 2001; Leinonen *et al*., 2008).

## 2. Materials and Methods

### 2.1 Study area and sampling design

The study area included five lake ecosystems (sites hereafter) in both Central and Northern Italy (Figure 1): Lake Chiusi (“CH”; 43° 03′ 22′′ N, 11° 57′ 56′′ E); Lake Massaciuccoli (“MA”, 43° 50′ 0′′ N, 10° 19′ 30′′ E); Mantua lakes system (“MN”, 45° 09′ 36″ N, 10° 47′ 48″ E); Lake Iseo, including the Torbiere del Sebino wetland (“IS”, 45° 43′ 00″ N, 10° 05′ 00″ E); lakes Pusiano and Annone (“PA”, 45° 48′ 40″ N, 9° 18′ 45″ E). They are phytogeographically comparable sites covering a wide environmental gradient, in terms of ecological conditions (i.e., trophic status), geographical features (i.e., size and watershed land cover), and plant community structure (i.e., macrophyte community types, or growth forms). The five lake systems differ in areal size, ranging from ∼3.8 km^2^ to 65.3 km^2^. In terms of trophic conditions, the lakes range from oligo-mesotrophic (Iseo) and mesotrophic (Pusiano), up to eutrophic (Annone) and even hypertrophic conditions (Chiusi, Massaciuccoli and Mantova).

**Figure 1.**
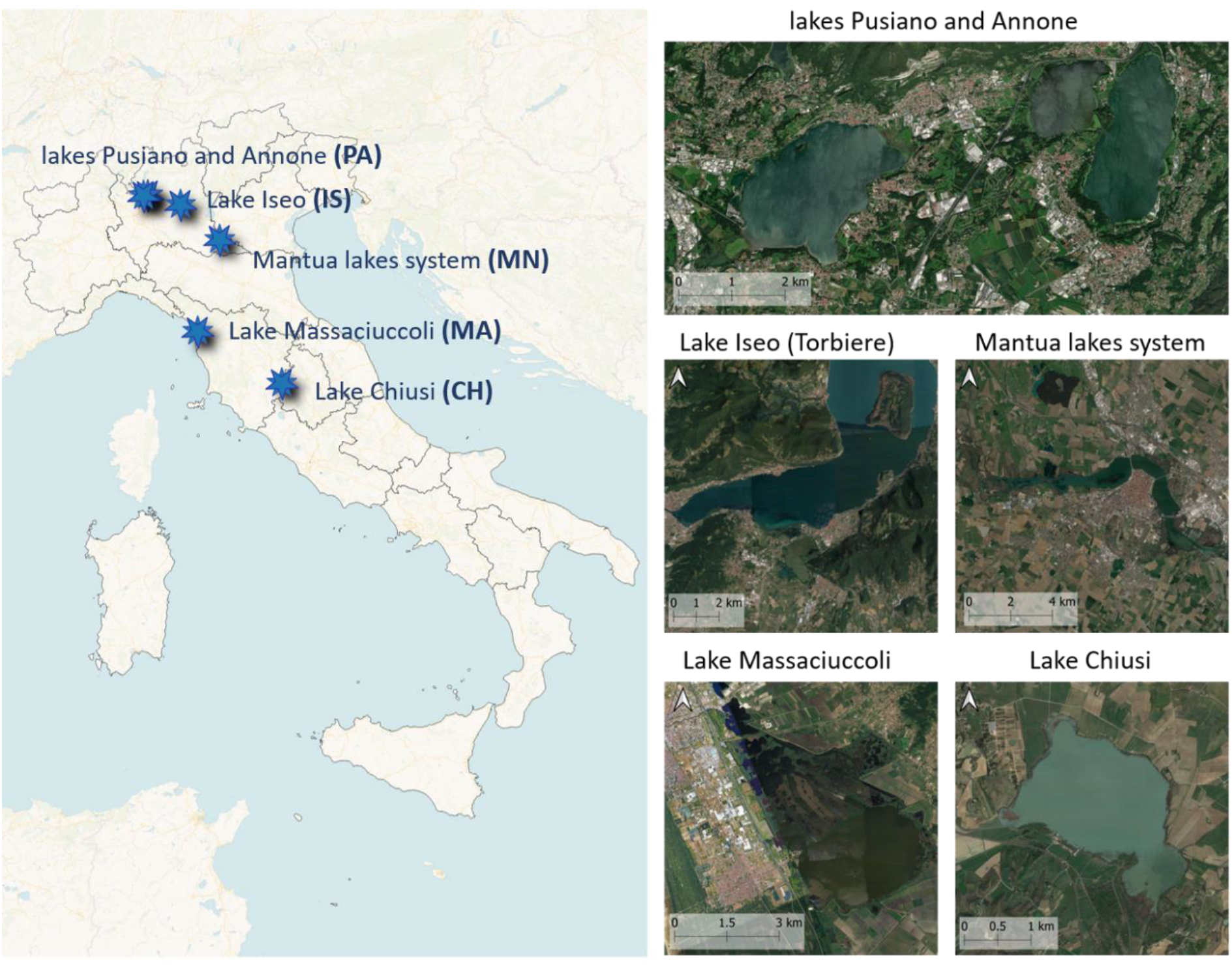
Geographic location of the study area with distribution of the sampling sites. Satellite images of each site (at different scales), taken from QGis’ Google Satellite Basemap, are shown on the right.

Depending on the lake area and the relative coverage of the target plant communities (*P. australis* and *N. lutea*), 10 m x 10 m plots were sampled in each site to be representative of the local ecological heterogeneity (*N. lutea* was not present in Massaciuccoli Lake) [Supplementary material Table S1]. Out of the total 78 sampled plots, 50 were dominated by *P. australis* and 28 by *N. lutea*. Eight leaves were randomly taken from each plot for the functional, spectral, or genetic analyses. When it was not possible to use the very same leaf for two or more analyses types (e.g., leaf traits and spectral reflectance), leaves from the same culms (for *P. australis*) or same rosettes (for *N. lutea*) were collected as matches.

### 2.2 Spectral data

Reflectance in the visible to shortwave infrared spectrum range (350 – 2500 nm) was measured from all the fresh leaves within seconds (maximum 1 minute) after cutting them from the plants. Measurements were done using a portable high resolution spectroradiometer (SR-3500, Spectral Evolution, Lawrence, USA), with a spectral resolution of 3 nm for wavelengths under 1000 nm, and < 8 nm up to 2500 nm. To minimize disturbance due to background reflection of transmitted light, leaves were placed on a black neoprene plate (reflectance factor < 5%) during spectra measurements. Leaf reflected radiance was measured with a contact probe with an internal light source (5 W) as an average of 10 scans, and finally calibrated to reflectance using as reference the contact probe readings taken over a Spectralon panel (Labsphere, North Sutton, USA; reflectance factor > 95%).

### 2.3 Functional traits

Five of the eight intact, well-developed leaves of *N. lutea* and *P. australis* were used to measure structural traits (LA, mm^2^; SLA, mm^2^/mg), while three were used to measure bio-chemical traits (Chl-a, μg/g; Chl - a/Chl-b). After gently cleaning water-saturated leaves of debris and epiphytes, they were scanned for measuring LA. Leaves were subsequently dried at 50°C until constant weight to quantify the dry weight (mg). LA was determined analyzing scanned images with the software ImageJ (Rasband, 1997, 2018), whereas SLA was calculated as the ratio between LA and dry weight. High values of SLA indicate an acquisitive behavior, implying lower investments in structural tissues for photosynthetic organs and a higher photosynthetic capacity *per* mass unit (Dalle Fratte *et al*., 2019b). Chlorophyll content was determined spectrophotometrically after 24-h extraction in 80% acetone (Wellburn, 1994).

Additional leaf traits were derived from the inversion of PROSPECT-D model starting from leaf reflectance. PROSPECT-D is a physical model based on radiative transfer theory that simulates leaf optical properties (reflectance and transmittance) in the spectrum domain ranging from 400 to 2500 nm, based on leaf biochemical constituents (chlorophylls, carotenoids, anthocyanins, water, and dry matter) and an anatomical structural parameter, termed N (Féret *et al*., 2017). PROSPECT-D was inverted through iterative optimization, minimizing the RMSE between measured and simulated reflectance with optimal spectral domain and configuration for each individual trait (Spatford *et al*., 2021), using the R package “prospect” (Féret and de Boissieu, 2022; Team RC, 2019). The four traits derived from PROSPECT-D model inversion output are: total chlorophylls (Chl_ab); dry matter content on area basis, or leaf mass per area (LMA); dry matter content on weight basis or leaf dry matter content (LDMC); and the mesophyll structure parameter, as a proxy for mesophyll complexity (Nmesophyll). As PROSPECT was designed and calibrated on terrestrial plant species, to increase the accuracy of prediction over aquatic plants, species-specific correction factors were applied to raw model inversion output for Chl_ab, LMA and LDMC, separately for *P. australis* and *N. lutea* samples, based on actual trait scores measured over a subset of sampled leaves (N = 238-394, varying with trait). Such recalibration reduced the relative error of modelled traits by 1 to 10 % depending on the trait, with nRMSE ranging from 6% (LDMC) to 12% (Chl_ab).

### 2.4. DNA extraction and AFLP protocol

A total of 400 and 206 samples (from five to eight leaves per plot) were analyzed for *P. australis* and *N. lutea*, respectively. Each leaf tissue sample was ground in a mortar with sterile sand. DNA extraction was carried out using the 2x cetyltrimethylammonium bromide (CTAB) protocol (Doyle and Doyle, 1990). Quality and quantity control of extracted DNA were performed using Bio-Photometer (Eppendorf, Germany). AFLP analysis was performed with minor modifications of previous studies using molecular tools (see Coppi *et al*., 2014 and references therein). Two combinations of primers were selected for analyses in both species: hex_EcoRI-CTA/MseI-ATG and fam_EcoRI-TAC/MseI-ATG for *P. australis* and hex_EcoRI-ACG/MseI-TTA and fam_EcoRI-CTA/MseI-CTC for *N. lutea*. AFLP profiles obtained by capillary electrophoresis were analyzed using GeneMarker v1.5 (SoftGenetics LLC, State College, PA, United States).

### 2.5. Analyses of genetic variation at sampling plot and site level

The average genetic diversity within a sampling plot (AGD) was computed as the probability that two homologous sites are different (Nei, 1987) by using Arlequin v2.000 (Schneider *et al*., 2000). The percentage composition of polymorphic bands was calculated as [(nfrag/ntotal)*100], where nfrag was AFLP loci detected for each sampling plot or site and ntotal the number of total detected bands for the primer pair.

As for the genetic structure, molecular variance analysis (AMOVA, Excoffier et al., 1992) was performed using Arlequin v2.000 (Schneider *et al*., 2000) to determine the distribution of total genetic variation at different hierarchical levels: i) within and among sampling plot, ii) within and among sites. The analyses were performed separately for the two hierarchical levels. Tests of the variance components and the percentage of total expressed variation were conducted to assess the statistical support for the different groups. Genetic distances between populations and sites were estimated by computing Slatkin’s linearized pairwise *F*_st_ values (Slatkin, 1995). Following Yang *et al*. (2016), loci under selection (outliers) were detected using BayeScan v2.01 to remove them from the calculation of neutral genetic differentiation. Outliers are loci that fall over a threshold value set on the logarithm of posterior odds values (LogPO), determined as in Foll (2012). The number of pilot runs was kept at 20, with a length of 10 000 iterations each (Coppi *et al*., 2018).

### 2.6. P_st_ - F_st_ comparison

Phenotypic variance (*P*_st_), estimated from functional leaf traits, were compared to *F*_st_ values using “Pstat” package (Silva and Silva, 2018) in R environment. *F*_st_ values for both species were extrapolated from the molecular variance analysis [see Supplementary Material Tables S2 and S3]. For each site, *F*_st_ values at population level were averaged and bootstrapped 95% confidence intervals were calculated using R. *P*_st_ values were determined with the bootstrap method (1,000 x) under a confidence level of 95% using “Pstat” package (Silva and Silva, 2018) [see Supplementary Material Tables S4].

*P*_st_ was calculated from between population (σ^2^_B_) and within population (σ^2^_w_) components of variance for each trait following Brommer’s (2011) expression:

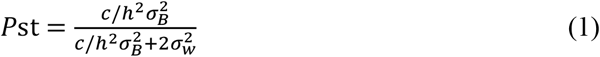

where c is the proportion of the total variance attributed to additive genetic effects between populations and h^2^ is the heritability *stricto sensu*. In this index, the c/h^2^ ratio quantifies the proportion of phenotypic differences observed between populations that can be attributed to additive genetic variance (Leinonen *et al*., 2008; Brommer, 2011). The problem of using *P*_st_ as an approximation of *Q*_st_ is mainly because values of c/h^2^ are unknown in natural populations (Pujol et al., 2008). Consequently, the starting point is the null assumption c/h^2^= 1 (i.e., c=h^2^). However, natural populations may be subject to genotype-environmental interactions and low values of c/h^2^ (i.e, c < h^2^) assumes a more important role of environmental factors in determining between-population variance than within-population variance. Therefore, the lower the critical c/h^2^ ratio is (c/h^2^ < 1) when *P*_st_ exceeds *F*_st_, the more likely it is that the trait is being shaped by selection (Brommer, 2011). Dealing with this issue, we tested three different c/h^2^ (0.5, 0.63, 1 [Supplementary material Table S4]) and we eventually assessed the strength of *P*_st_ – *F*_st_ comparisons for the more conservative case (c/h^2^ = 0.5).

Cohen’s d was used to calculate the effect size for *P*_st_ - *F*_st_ differences:

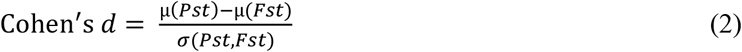

where μ(*P*_st_) is the *P*_st_ value computed from “Pstat”, μ(*F*_st_) is the average *F*_st_ score, and σ(*P*_st_, *F*_st_) is the pooled standard deviation derived as the sum of σ(*P*_st_) and σ(*F*_st_). to derive pooled standard deviation σ(*P*_st_) is estimated using the lower bound of 95% confidence intervals computed from “Pstat” as 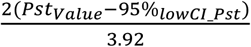, and σ(*F*_st_) is estimated using the 95% confidence intervals of *F*_st_ average as 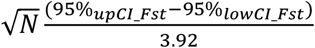.

As a first interpretation, we categorized the continuous range of Cohen’s *d* scores into three situations, following the scale proposed by Sawilowsky (2009), denoting dominant mechanisms underlying intraspecific diversity in terms of *P*_st_ - *F*_st_, such as: stabilizing selection (SS), when d ≤ -0.8; directional or divergent selection (DS), when d ≥ 0.8; and an intermediate situation indicating neutral selection (NS), when -0.8 < d < 0.8.

To investigate whether natural selection dominance (DS) is driven by directional or divergent evolution patterns, the distribution of trait values and AFLP mismatches were assessed in both species and within each site [see Supplementary Material Fig. S2 and Fig. S5, respectively]. Mismatch distributions (i.e., the distribution of pairwise differences among haplotypes; Rogers and Harpending, 1992) were calculated using Arlequin v2.000 software (Schneider *et al*., 2000). Unimodal distribution was interpreted as the effect of directional selection, while bimodal to multimodal distribution of pairwise differences or measured traits was interpreted as the effect of divergent selection (Choudhuri, 2014).

To determine whether *P*_st_ – *F*_st_ comparisons are biased by the choice of markers (Edelaar *et al*., 2011), the relationship between *P*_st_ and *F*_st_ difference and average genetic diversity over loci at site level was investigated and tested using a linear regression [see Supplementary Material Fig. S1].

## 3. Results

### 3.1. Functional traits

Overall, 250 and 130 leaves of *P. australis* and *N. lutea*, respectively, were analyzed for structural traits, and 150 and 84 leaves for bio-chemical traits. In *P. australis*, the median values of traits ranged as follows: LA from 6525.4 (IS) to 10536.0 mm^2^ (MN), SLA from 12.1 (MN) to 13.9 mm^2^ mg^-1^ (MA), Chl-a from 2021.4 (CH) to 3069.2 μg g^-1^ (MN), and Chl-a/Chl-b from 4.0 (CH) to 4.7 μg g^-1^ (PA). As for *N. lutea*, LA ranged from 56748.9 (IS) to 80657.0 mm^2^ (PA), SLA from 6.1 (PA) to 9.1 mm^2^ mg^-1^ (CH), Chl-a from 693.3 (PA) to 934.9 μg g^-1^ (IS), and Chl-a/Chl-b from 3.4 (CH) to 4.4 μg g^-1^ (IS). Individual measures and distributions are available in Supplementary Material [Table S5 and Fig. S2, respectively].

Concerning the PROSPECT-D derived traits, 400 and 224 leaves were analyzed for *P. australis* and *N. lutea*, respectively. In *P. australis*, the median values of traits ranged as follows: Chl_ab from 51.8 (CH) to 56.2 μg cm^-2^ (PA) and LMA from 72.9 g m^-2^ (MA) to 81.5 g m^-2^ (IS), LDMC ranged from 0.40 (MA) to 0.45 g g^-1^ (IS), and Nmesophyll from 1.41 (MA) to 1.57 (PA). As for *N. lutea*, Chl_ab ranged from 33.1 (CH) to 40.8 μg cm^-2^ (PA) and LMA from 76.8 (CH) to 108.2 g m^-2^ (PA), LDMC ranged from 0.16 (CH) to 0.19 g g-1 (PA), and Nmesophyll from 1.57 (CH) to 2.15 (PA). Individual measures and distributions are available in Supplementary Material [Table S5 and Fig. S2, respectively].

The intraspecific leaf reflectance variability and the comparison between leaf traits measured and estimated from PROSPECT-D inversion are shown in Supplementary Material [Fig. S3 and S4, respectively].

### 3.2. Amplified Fragment Length Polymorphisms Analysis

#### Phragmites australis

The AFLP analysis was successfully performed on 395 samples. The selected combinations of primers produced a total of 341 loci, 130 for the combination hex_EcoRI-CTA/MseI-ATG and 211 for that of fam_EcoRI-TAC/MseI-ATG. Within sampling plots, the percentage of polymorphic loci (PPL) ranged from a maximum of 87.7% (MN14) down to a minimum of 29.6% (CH08). Within each site, the average percentage of polymorphic loci varied from 97.4% in Mantova to 76.8% in Chiusi. The AGD levels varied from 0.086 (CH02) to 0.353 (MA08) within sampling plot and from 0.176 to 0.321 (Chiusi and Mantova respectively) at site level. Genetic diversity values both at sampling plot and site level are shown in Supplementary Material [Table S2]. Regarding AMOVA, the greatest percentage of the total genetic variation occurred within sampling plots (66.9%), rather than among sampling plots (33.1%) (Table 1a). The same can be observed across sites, where genetic differentiation within sites (79.3%) was higher than among sites (20.7%) (Table 1b). The BayeScan analysis identified one outlier locus that had a posterior probability greater than 0.78 (at a threshold of log10 PO > 0.5).

**Table 1.** The partition of genetic variance (AMOVA) for P. australis was represented at two different hierarchical levels: a) within and among sampling plot, b) within and among sites. Tables show degrees of freedom (d.f.), sum of squared deviations, variance component estimates, percentages of total variance contributed by each component, and probability of obtaining a more extreme component estimate by chance alone (p). P-values were estimated with 999 permutations.

| **a)** |  |  |  |  |  |
| --- | --- | --- | --- | --- | --- |
| Source of variation | d.f. | Sum of squares | Variance Components | Percentage of variation | P-values |
| Among sampling plots | 49 | 9200.153 | 18.92043 | 33.07 | <0.0001 |
| Within sampling plots | 345 | 13210.464 | 38.2912 | 66.93 | <0.0001 |
| Total | 394 | 22410.618 | 57.21164 | 100 |  |
| **b)** |  |  |  |  |  |
| Source of variation | d.f. | Sum of squares | Variance Components | Percentage of variation | P-values |
| Among sites | 4 | 3955.551 | 12.32683 | 20.67 | <0.0001 |
| Within sites | 390 | 18455.067 | 47.32069 | 79.33 | <0.0001 |
| Total | 394 | 22410.618 | 59.64751 | 100 |  |

#### Nuphar lutea

The AFLP analysis was successfully performed on 203 samples. The selected combinations of primers produced a total of 191 loci, 94 for the combination hex_EcoRI-ACG/MseI-TTA and 97 for that of fam_EcoRI-CTA/MseI-CTC. Within sampling plots, the percentage of polymorphic loci (PPL) ranged from a maximum of 92.7% (IS06) down to a minimum of 35.6% (CH17). Within each site, the average percentage of polymorphic loci varied from 86.4% for Iseo to 77.5% for Pusiano - Annone. The AGD levels varied from 0.098 (CH05) to 0.287 (MN33) within sampling plots and from 0.212 to 0.267 (Chiusi and Iseo, respectively) at the site level. Genetic diversity values both at intra- and inter-site level are shown in Supplementary Material [Table S3]. Regarding AMOVA, the percentage of the total genetic variation was nearly equal within (53.5%;) and among sampling plots (46.5%) (Table 2a) and within (54.1%) and among (45.9%) sites (Table 2b). BayeScan analysis did not detect any outlier loci.

**Table 2.**
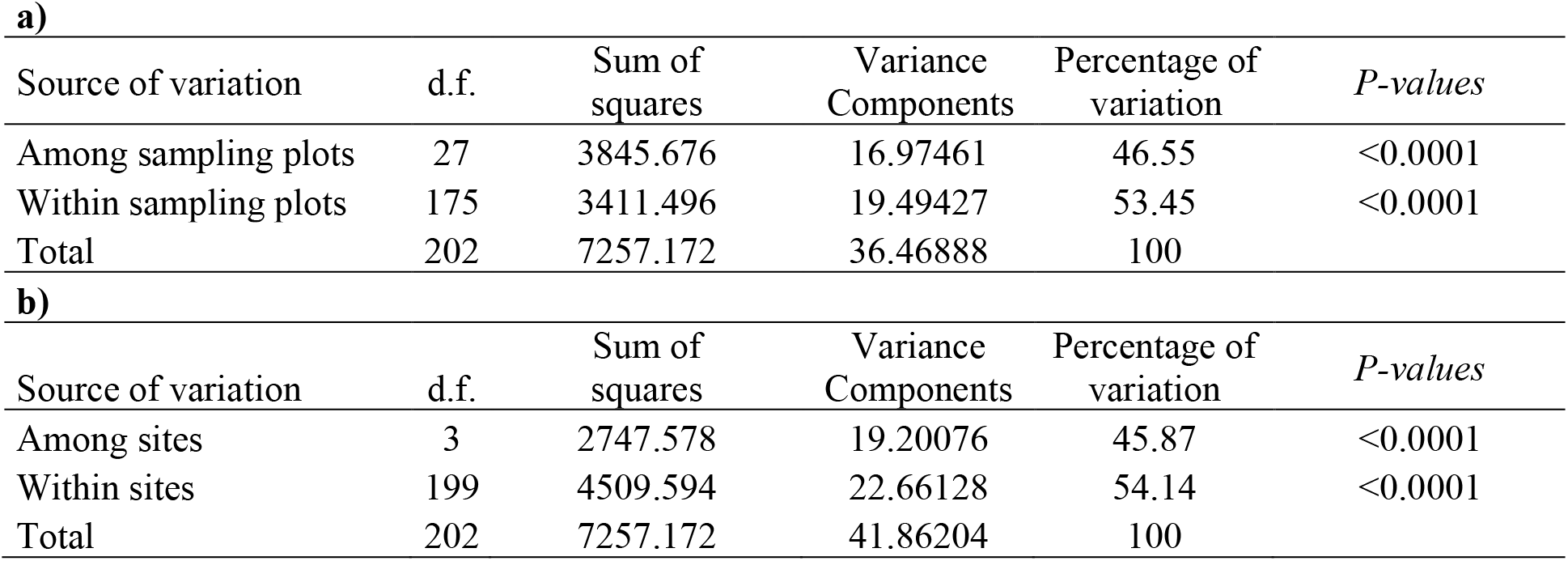
The partition of genetic variance (AMOVA) for N. lutea was organized at two different hierarchical levels: a) within and among sampling plot, b) within and among sites. Tables show degrees of freedom (d.f.), sum of squared deviations, variance component estimates, percentages of total variance contributed by each component, and probability of obtaining a more extreme component estimate by chance alone (p). P-values were estimated with 999 permutations.

| **a)** |  |  |  |  |  |
| --- | --- | --- | --- | --- | --- |
| Source of variation | d.f. | Sum of squares | Variance Components | Percentage of variation | P-values |
| Among sampling plots | 27 | 3845.676 | 16.97461 | 46.55 | <0.0001 |
| Within sampling plots | 175 | 3411.496 | 19.49427 | 53.45 | <0.0001 |
| Total | 202 | 7257.172 | 36.46888 | 100 |  |
| **b)** |  |  |  |  |  |
| Source of variation | d.f. | Sum of squares | Variance Components | Percentage of variation | P-values |
| Among sites | 3 | 2747.578 | 19.20076 | 45.87 | <0.0001 |
| Within sites | 199 | 4509.594 | 22.66128 | 54.14 | <0.0001 |
| Total | 202 | 7257.172 | 41.86204 | 100 |  |

### 3.3. P_st_ - F_st_ comparison

The values of effect size for *P*_st_ - *F*_st_ comparison are summarized in Table 3. The distributions of trait values [Supplementary Material Fig. S2] and mismatches [Supplementary Material Fig S5] were generally unimodal, suggesting that, in the case of *P*_st_ > *F*_st_ by a reasonable extent (Cohen’s *d* > 0.8), directional selection could be the main force leading to population differentiation. Some exceptions were evident for Nmesophyll in PA for *P. australis*, and for LA in CH, IS and MN or LMA in IS for *N. lutea*, indicating that in a minority of situations (specific sites and traits) the action of divergent selection cannot be excluded.

**Table 3.** Results of Pst - Fst comparison in terms of effect size (Cohen’s d) for each trait within sites (CH= Chiusi; IS= Iseo; MA= Massaciuccoli; MN= Mantova; PA= Pusiano-Annone). The higher and positive Cohen’s d scores are (Pst > Fst), the more evidently phenotypic differentiation of a specific trait is likely driven by directional or divergent selection (DS, highlighted in blue). Conversely, the higher and negative Cohen’s d scores are (Pst < Fst), the more evidently phenotypic differentiation of a specific trait is likely driven by stabilizing selection (SS, highlighted in red). In the middle, Cohen’s d scores waggling around zero indicate that neutral divergence (NS, not highlighted) cannot be excluded as a possible cause of phenotypic diversity (Pst = Fst).

| *P. australis* |  |  |  |  |  | *N. lutea* |  |  |  |  |
| --- | --- | --- | --- | --- | --- | --- | --- | --- | --- | --- |
| Trait | CH | IS | MA | MN | PA | Trait | CH | IS | MN | PA |
| Chl_ab $\mu\text{g cm}^{-2}$ | 0.74 | -0.68 | 3.02 | 2.37 | 1.48 | Chl_ab $\mu\text{g cm}^{-2}$ | 2.15 | 0.73 | 1.35 | -1.77 |
| LMA $\text{g m}^{-2}$ | -1.7 | 0.69 | 1.98 | 2.29 | 2.43 | LMA $\text{g m}^{-2}$ | 0.67 | 1.92 | -0.29 | -1.02 |
| LDMC $\text{g g}^{-1}$ | -0.24 | 2.41 | 0.24 | 2.29 | 3.38 | LDMC $\text{g g}^{-1}$ | -0.89 | 1.49 | -1.09 | -0.38 |
| Nmesophyll | 1.72 | 0.54 | 3.43 | 3.74 | 7.93 | Nmesophyll | 0.42 | -0.35 | 0.93 | 0.85 |
| Leaf.area.tot $\text{mm}^2$ | 1.84 | 1.9 | 2.5 | 0.96 | 1.34 | Leaf.area.tot $\text{mm}^2$ | 2.61 | 5.21 | 3.05 | 0 |
| Leaf.SLA.$\text{mm}^2 \text{mg}^{-1}$ | -1.26 | -0.78 | 1.87 | 1.08 | 0.19 | Leaf.SLA.$\text{mm}^2 \text{mg}^{-1}$ | -0.54 | 0.25 | 9.05 | -0.48 |
| Chl.a $\mu\text{g/g}$ | 1.84 | 0.1 | 1.14 | 0.81 | 1.63 | Chl.a $\mu\text{g g}^{-1}$ | -0.75 | -0.35 | 2.06 | -1.44 |
| Chl.a/Chl.b $\mu\text{g g}^{-1}$ | -2.14 | 0.43 | -0.43 | 0.44 | -1.8 | Chl.a/Chl.b $\mu\text{g g}^{-1}$ | 0.35 | -0.69 | -1.16 | -3.84 |

In *P. australis* populations, directional selection is very evident as the main driver of phenotypic differentiation (Cohen’s *d* >2, i.e., *P*_st_ is higher than *F*_st_ by at least two times the pooled standard deviation) in MA, MN, and PA sites across many traits, i.e., 4 out of 8 traits in the former two sites and 2 out of 8 in the latter (plus Nmesophyll possibly driven by divergent selection). The patterns are generally leaning toward neutral divergence for IS populations (except for LDMC and LA, driven by directional selection with Cohen’s *d* > 1.9), and a mixture of directional (Nmesophyll, LA, Chl-a) and stabilizing (LMA, SLA, Chl-a/Chl-b) selection in CH populations. In addition, there was a significant, yet weak relationship between AGD values and the *P*_*st*_ *– F*_*st*_ for LMA, suggesting a possible bias of *P*_*st*_ *– F*_*st*_ for this trait.

As for *N. lutea*, stabilizing selection (marked by large, negative Cohen’s *d* of *P*_st_ - *F*_st_ difference) appears to be the dominant driver of phenotypic differentiation in PA populations (especially marked for pigments-related traits, Cohen’s *d* < -1.4). The populations in CH showed a mixed pattern, i.e., the relative dominance of neutral selection for several traits (5 out of 8), with the exceptions of LA and Chl-a/Chl-b (apparently driven by divergent and directional selection, respectively), and LDMC (SS-driven). Similarly to what happens for *P. australis*, the dominance of neutral selection is observed for *N. lutea* populations in IS (5 out of 8 traits), while LA and LMA variation appears to be driven by divergent selection, and LDMC by directional selection. Directional selection seems to be the main force driving most of the FTs’ differentiation in MN populations (4 out of 8 traits), in particular for SLA, scoring average *P*_st_ values higher than *F*_st_ by nine times the pooled standard deviation, while LA appears to be driven by divergent selection, from the bimodal distribution shown in Fig. S2a [Supplementary Material].

## 4. Discussion

In all the five investigated sites, the genetic structure of *P. australis* agrees with other Italian and European conspecific populations, showing a higher genetic variation within rather than among plots and sites (Lambertini *et al*., 2008; Coppi *et al*., 2018). *N. lutea*, on the other hand, exhibited a more complex genetic structure than expected at site level, possibly due to the combined effect of genetic isolation and its mixed mating system. Both species showed wide variability in leaf functional traits within and across sites, confirming their high plasticity (Eller and Brix, 2012; Kordyum and Klimencko, 2013; Guo *et al*., 2016; Dalla Vecchia and Bolpagni, 2022). As for evolutionary drivers, *P*_st_ - *F*_st_ comparisons showed an overall tendency to directional selection for *P. australis* and a more complex pattern for *N. lutea*.

### 4.1. Genetic structure and traits variation

The mean genetic diversity indices at both sampling plot and site levels showed no obvious signs of gene erosion for *P. australis*, apart from populations in Chiusi, characterized by lower genetic diversity than other sites. AGD and PPL values were in line with those observed for other conspecific populations in Italy and Europe (Lambertini *et al*., 2008; Coppi *et al*., 2018). Consistently, most of the genetic variation resided within rather than among sampling plots and sites, confirming previous studies on phylogeographic relationships in common reed stands on local to narrow range scales (Lambertini *et al*., 2008; Qiu *et al*., 2016; Coppi *et al*., 2018). Indeed, the high local genetic diversity observed for *P. australis* agrees with previous evidence, and it seems directly connected with the ecological and physiological characteristics of the species (Gao *et al*., 2012; Richards *et al*., 2012). As an example, invasiveness of common reed genotypes in North America has been attributed the capacity of this species to combine sexual reproduction and vegetative propagation (McCormick *et al*., 2010; Albert *et al*., 2015).

As for *N. lutea*, the two previous works on this topic showed a tendency towards greater genetic diversity within than among populations (Fér and Hroudova, 2008; Vyšniauskiene *et al*., 2020). Our results, covering relatively distant sites (up to 370 km), show a partition of genetic variation nearly equal within and among both sampling plots and sites, thus revealing the existence of complex genetic structure characterizing the targeted *N. lutea* populations. The ability of *N. lutea* to shift its reproduction strategy from out- or in-breeding (Ervik *et al*., 1995; Lippok and Renner, 1997; Padgett, 2007) to the vegetative and vice versa strongly affects the dynamics of populations and, consequently, their genetic structure. Merging the findings of previous works (e.g., Fér and Hroudova, 2008; Padgett, 2007) with our genetic outcomes – in particular the complete lack of clonal profiles – allows us to outline an interpretative framework for the establishment and expansion of *N. lutea* populations in lentic ecosystems. During a first “establishment step” (i), new germplasm of mainly generative origin arrives from nearby populations, entailing high genetic diversity within the new population and low genetic distance from donor populations; ii) during a second “expansion wave”, vegetative reproduction leads to competition among genotypes, favoring individuals well adapted to local conditions that will tend to outcompete new migrants, and results in lower genetic diversity within the population and higher genetic distance from donor populations. Genetic drift may continue as long as the filtering of genotypes occurs, or new, competitive migrants succeed in establishing. However, it should not be excluded that genetic differentiation of the new population from the donor populations may also be increased due to sexual reproduction of the filtered genotypes.

In this context, the mixed reproductive strategy could play a role in how both *P. australis* and *N. lutea* respond to environmental variation. However, it is crucial to note that phenotypic variation can also promote adaptive evolutionary responses even if induced by the environment and not strictly controlled by genetics (Waddington, 1952, 1953; West-Eberhard, 2003). Regardless of the pre-eminent evolutive process, evaluated by the *P*_*st*_ *- F*_*st*_ comparisons, high rates of leaf traits variation were observed across sites, suggesting a high phenotypic plasticity both in structural and biochemical traits. These results reinforce the outcomes by Dalla Vecchia and Bolpagni (2022), who found high rates of intraspecific variability for *N. lutea* at the micro-scale (Lake Chiusi), as well as strong links between traits variation and environmental drivers, especially for leaf size (LA and dry weight).

### 4.2. P_st_ - F_st_ comparison

At present, local adaptation capabilities for *P. australis* and *N. lutea* have been studied from either a phenotypic (Vretare *et al*., 2001; Henriot *et al*., 2019; Ren *et al*., 2020) or a genetic point of view (Fér and Hroudova, 2008; Coppi *et al*., 2018; Vyšniauskiene *et al*., 2020 Lambertini *et al*., 2020; Naugžemys *et al*., 2021). To our knowledge, only one study has jointly investigated the complex relationships between population-based functional trait variation, genetic diversity, and environmental heterogeneity, i.e., the work of Wani *et al*., (2020) focusing on *P. australis*. For *P. australis*, the outcomes of our *P*_st_ - *F*_st_ comparison indicated that for a majority of sites (MA, MN, and PA) most of the observed FTs variation was affected by directional selection, thus led by genetic adaptation.

In target *P. australis* populations, directional selection tends to move the traits mean towards the optimum for that environment, increasing the adaptability of individuals, thus confirming the observations of Wani and colleagues (2020) for the invasive haplotypes, and indicating the importance of the investigation *P*_st_ - *F*_st_ comparisons at a fine local scale (i.e., among sites within geographic regions). In this context, the proportion of variation in the phenotype due to neutral genetic differentiation approaches zero and the remaining phenotypic variation is from either environmental or adaptive genetic variation. This result, coupled with the genetic outcomes, confirms that the colonization strategy of *P. australis* implies a prevalence of patches of individuals with high genetic diversity, well adapted to local conditions, across most of the studied sites. It is hypothesized that the isolation of patches of individuals and populations showing high local adaptation could be the effect of the prevalence of vegetative over generative reproduction (Koppitz and Kuehl, 2000; Alvarez *et al*., 2005; Lebedeva *et al*., 2020). Exceptions to this pattern are the CH and IS populations, where traits variability is mainly shaped respectively by directional and stabilizing selection (DS and SS), and by directional and neutral divergence (DS and NS). Our results show that Chiusi and Iseo sites host lower genetic diversity and higher genetic differentiation for neutral markers than other sites, suggesting that patches of common reed are undergoing genetic drift at the micro-local scale, which in turn promotes SS on selected leaf traits. Despite this, in these sites the FTs did not show substantial differences compared to the others, except for the lowest values for Chl-a and the highest average value regarding the Chl-a and -b ratio in CH, the hypertrophic site.

On the other hand, in *N. lutea* the *P*_st_ - *F*_st_ comparison showed differently mixed patterns, ranging from SS dominance for populations in PA to different balances of stabilizing, neutral, and directional or divergent selection in the other sites. As the rate of plant adaptation to the environment depends on how close the plastic phenotype is to the environmental optimum (Price *et al*., 2003), when plasticity matches ideal conditions, the population should undergo SS without subsequent genetic differentiation; this is the case for *N. lutea* in PA across all the leaf pigments and LMA, contrasted by Nmesophyll (directional selection-affected). Instead, when traits tend to be far from the environmental optimum, the population should be subjected to directional selection. However, where divergent phenotypic patterns are present across populations at the same site, divergent selection may be the main driver, as in the case of LA in CH, IS and MN. These evolutionary trajectories seem to only partially reflect the trophic gradient among sites, with PA showing intermediate quality status (meso-eutrophic). Conversely, IS and the MN-CH pair, where the leading process is represented by DS, show oligo-meso- and hypertrophic conditions, respectively.

The use of *P*_st_ - *F*_st_ comparison allowed us to infer the relative role of genetic drift and natural selection on the diversification of their leaf traits across different lake systems. Our results showed an overall tendency to directional selection for *P. australis*, with the exclusion of the Chiusi site where stabilizing selection drives the variability of three out of eight traits. On the other hand, *N. lutea* showed a more complex pattern of phenotypic differentiation drivers, with a mixture of stabilizing, neutral and directional (divergent) selection in most of the investigated sites. These outcomes suggest the existence of species-specific behaviors in the diversification of leaf traits for both macrophytes, with marked differences between the two species in the PA site (Table 3). Conversely, populations of both species in IS showed similar differentiation patterns (dominated by neutral selection), indicating a sensitive effect of site conditions in affecting their functional plasticity. In this respect, it will be necessary to direct future analyses towards a better understanding of the implications of ecosystem quality status in driving local adaptation processes and functional responses of macrophytes, in order to define the extent and limits of their plastic adaptation across a wider range of conditions.

The sampling and characterization of *P. australis* and *N. lutea* populations inhabiting the same sites allowed us, for the first time, to perform a joint analysis of the genetic structure of both species. This is a fundamental point for understanding the relative roles of genetic drift and natural selection on the diversification of phenotypic traits within the habitat of two keystone macrophyte species. As the relative contribution of selection, genetics, and plasticity to environmental adaptation appears to be strictly determined by the ecological context, further studies are needed to corroborate these initial findings and to deepen our understanding of the relative role of natural selection in the diversification of two target species and how this relates to eco-geographical variables.

## Supplementary Materials

Supplementary data are available online at https://…… and consist of the following Tables and Figures: Tab. S1: Schematic representation of the experimental design. Tab. S2: Genetic diversity values of *P. australis*. Tab S3: Genetic diversity values of *N. lutea*. Tab. S4: *P*_st_ and *F*_st_ values for each trait at site level. Tab. S5: Functional traits values for each species. Fig. S1: relationship between the difference between Pst and Fst and within-site avarage genetic diversity over loci for every investigated trait. Fig. S2: Violin plots showing the variability of *P. australis* and *N. lutea* functional traits. Fig. S3: Intraspecific leaf reflectance variability. Fig. S4: Comparison between leaf traits estimated from PROSPECT-D inversion with or without applying any correction factor. Fig. S5: Mismatch distributions of *P. australis* and *N. lutea* within each site.

## Acknowledgements

This work was supported by the project “macroDIVERSITY”, funded by the Ministry of Education, University and Research, PRIN 2017 [grant number 2017CTH94H]. ADV has benefited from the equipment and framework of the COMP-HUB Initiative, funded by the ‘Departments of Excellence’ program of the Italian Ministry for Education, University and Research (MIUR, 2018-2022).

## Author Contributions

Conceptualization: AC, PV. Developing methods: MBC, AC, RB, PV. Data analysis: MBC, AC, ADV, PV.

## References

Albert A, Brisson J, Belzile F, Turgeon J, Lavoie C. 2015. Strategies for a successful plant invasion: the reproduction of Phragmites australis in north-eastern North America. Journal of Ecology 103: 1529–1537.

Alvarez MG, Tron F, Mauchamp A. 2005. Sexual versus asexual colonization by Phragmites australis: 25-year reed dynamics in a Mediterranean marsh, southern France. Wetlands 25: 639–647.

Andrews KR, Good JM, Miller MR, Luikart G, Hohenlohe PA. 2016. Harnessing the power of RADseq for ecological and evolutionary genomics. Nature Reviews Genetics 17: 81–92.

Bolnick DI, Amarasekare P, Araújo MS, et al. 2011. Why intraspecific trait variation matters in community ecology. Trends in ecology & evolution 264: 183–192.

Bolnick DI, Svanbäck R, Fordyce JA, et al. 2003. The ecology of individuals: incidence and implications of individual specialization. The American Naturalist 161: 1–28.

Brommer JE. 2011. Whither P_ST_? The approximation of Q_ST_ by P_ST_ in evolutionary and conservation biology. Journal of evolutionary biology 24: 1160–1168.

Castellani MB, Coppi A., Bolpagni R, et al., 2023. Assessing the haplotype and spectro-functional traits interactions to explore the intraspecific diversity of common reed in Central Italy. Hydrobiologia 850: 775–791.

Chapuis E, Martin G, Goudet, J. 2008. Effects of selection and drift on G matrix evolution in a heterogeneous environment: a multivariate Q_st_–F_st_ test with the freshwater snail Galba truncatula. Genetics 180: 2151–2161.

Chave J. 2013. The problem of pattern and scale in ecology: what have we learned in 20 years? Ecology letters 16: 4–16.

Choudhuri S. 2014. Bioinformatics for beginners: genes, genomes, molecular evolution, databases and analytical tools. Elsevier.

Coppi A, Lastrucci L, Cappelletti D, et al. 2018. AFLP approach reveals variability in Phragmites australis: implications for its die-back and evidence for genotoxic effects. Frontiers in plant science 9: 386.

Coppi A, Cecchi L, Mengoni A, Pustahija F, Tomović G, Selvi F. 2014. Low genetic diversity and contrasting patterns of differentiation in the two monotypic genera Halacsya and Paramoltkia Boraginaceae endemic to the Balkan serpentines. Flora-Morphology, Distribution, Functional Ecology of Plants 2091: 5–14.

Da Silva SB, Da Silva A. 2018. Pstat: An R Package to Assess Population Differentiation in Phenotypic Traits. R J 101: 447.

Dalla Vecchia A, Bolpagni R. 2022. The importance of being petioled: leaf traits and resource-use strategies in Nuphar lutea. Hydrobiologia 849: 3801–3812.

Dalla Vecchia A, Villa P, Bolpagni R. 2020. Functional traits in macrophyte studies: Current trends and future research agenda. Aquatic Botany 167: 103290.

Dalle Fratte M, Bolpagni R, Brusa G, et al. 2019a. Alien plant species invade by occupying similar functional spaces to native species. Flora 257: 151419.

Dalle Fratte M, Brusa G, Pierce S, Zanzottera M, Cerabolini BEL. 2019b. Plant trait variation along environmental indicators to infer global change impacts. Flora 254:113–121.

Díaz S, Kattge J, Cornelissen JH, et al. 2016. The global spectrum of plant form and function. Nature 529: 167–171.

Doyle JJ, Doyle JL. 1990. Isolation of plant DNA from fresh tissue. Focus 12: 13–15.

Edelaar, PIM, Burraco, P, Gomez-Mestre, IVAN. 2011. Comparisons between QST and FST—how wrong have we been? Molecular Ecology, 20(23), 4830–4839.

Eid EM, Shaltout KH, Al-Sodany YM, Haroun SA, Jensen K. 2021. A comparison of the functional traits of Phragmites australis in Lake Burullus a Ramsar site in Egypt: Young vs old populations over the nutrient availability gradient. Ecological Engineering 166: 106244.

Eller F, Brix H. 2012. Different genotypes of Phragmites australis show distinct phenotypic plasticity in response to nutrient availability and temperature. Aquatic Botany 103: 89–97.

Ervik F, Renner S, Johanson KA. 1995. Breeding system and pollination of Nuphar luteum L Smith Nymphaeaceae in Norway. Flora 1902: 109–113.

Excoffier L, Smouse PE, Quattro J. 1992. Analysis of molecular variance inferred from metric distances among DNA haplotypes: application to human mitochondrial DNA restriction data. Genetics 1312: 479–491.

Fér T, Hroudova Z. 2008. Detecting dispersal of Nuphar lutea in river corridors using microsatellite markers. Freshwater Biology. 537: 1409–1422.

Féret JB, de Boissieu F. 2022. prospect: PROSPECT leaf radiative transfer model and inversion routines R package version 123.

Féret JB, Gitelson AA, Noble SD, Jacquemoud S. 2017. PROSPECT-D: Towards modeling leaf optical properties through a complete lifecycle. Remote Sensing of Environment 193: 204–215.

Gao L, Tang S, Zhuge, L, et al. 2012. Spatial genetic structure in natural populations of Phragmites australis in a mosaic of saline habitats in the Yellow River Delta, China

Guo WY, Lambertini C, Guo X, Li X-Z, Eller F, Brix H. 2016. Phenotypic traits of the Mediterranean Phragmites australis M1 lineage: differences between the native and introduced ranges. Biological Invasions 18: 2551–2561

Henriot CP, Cuenot Q, Levrey LH, et al. 2019. Relationships between key functional traits of the waterlily Nuphar lutea and wetland nutrient content. PeerJ 7: e7861.

Hughes AR, Inouye BD, Johnson MT, Underwood N, Vellend M. 2008. Ecological consequences of genetic diversity. Ecology letters 116: 609–623.

Karstens S, Jurasinski G, Glatzel S, Buczko U. 2016. Dynamics of surface elevation and microtopography in different zones of a coastal Phragmites wetland. Ecological Engineering 94: 152–163.

Kettenring KM, McCormick MK, Baron HM, Whigham DF. 2011. Mechanisms of Phragmites australis invasion: feedbacks among genetic diversity, nutrients, and sexual reproduction. Journal of Applied Ecology 485: 1305–1313.

Kiviat E. 2013. Ecosystem services of Phragmites in North America with emphasis on habitat functions. AoB plants 5.

Koppitz H, Kühl H. 2000. To the importance of genetic diversity of Phragmites australis in the development of reed stands. Wetlands Ecology and Management 86: 403–414.

Kordyum E, Klimenko E. 2013. Chloroplast ultrastructure and chlorophyll performance in the leaves of heterophyllous Nuphar lutea (L.) Smith. plants. Aquatic Botany 110: 84–91.

Lambertini C, Guo WY, Ye S, Eller F, Guo X, Li XZ, Brix H. 2020. Phylogenetic diversity shapes salt tolerance in Phragmites australis estuarine populations in East China. Scientific reports 101: 1–12.

Lambertini C, Gustafsson MH, Frydenberg J, Speranza M, Brix H. 2008. Genetic diversity patterns in Phragmites australis at the population, regional and continental scales. Aquatic Botany 882: 160–170.

Lebedeva OA, Belyakov EA, Lapirov AG. 2020. Reproductive potential of yellow water-lily Nuphar lutea in the conditions of lake ecosystems. Biosystems diversity 281: 60–67.

Leinonen T, McCairns RJ, O’hara RB, Merilä J. 2013. QST–FST comparisons: evolutionary and ecological insights from genomic heterogeneity. Nature Reviews Genetics 143: 179–190.

Leinonen T, O’hara RB, Cano JM, Merilä J. 2008. Comparative studies of quantitative trait and neutral marker divergence: a meta-analysis. Journal of evolutionary biology 211: 1–17.

Lenssen JP, Van Kleunen M, Fischer M, De Kroon H. 2004. Local adaptation of the clonal plant Ranunculus reptans to flooding along a small-scale gradient. Journal of Ecology 696–706.

Lippok B, Renner SS. 1997. Pollination of Nuphar (Nymphaeaceae) in Europe: Flies and bees rather than Donacia beetles. Plant systematics and evolution 2073: 273–283.

Maltchik L, de Oliveira GR, Rolon AS, Stenert C. 2005. Diversity and stability of aquatic macrophyte community in three shallow lakes associated to a floodplain system in the south of Brazil. Interciencia 303: 166–170.

Marin S, Gibert A, Archambeau J, Bonhomme V, Lascoste M, Pujol B. 2020. Potential adaptive divergence between subspecies and populations of snapdragon plants inferred from QST–FST comparisons. Molecular Ecology 29: 3010–3021.

Martin G, Chapuis E, Goudet J. 2008. Multivariate Q st–F st comparisons: a neutrality test for the evolution of the G matrix in structured populations. Genetics 1804: 2135–2149.

McCormick MK, Kettenring KM, Baron HM, Whigham DF. 2010. Extent and reproductive mechanisms of Phragmites australis spread in brackish wetlands in Chesapeake Bay, Maryland USA. Wetlands 301: 67–74.

Merilä J, Crnokrak P. 2001. Comparison of genetic differentiation at marker loci and quantitative traits. Journal of Evolutionary Biology 146: 892–903.

Mimura M, Yahara T, Faith DP, et al. 2017. Understanding and monitoring the consequences of human impacts on intraspecific variation. Evolutionary applications 102: 121–139.

Moran EV, Ormond RA. 2015. Simulating the interacting effects of intraspecific variation, disturbance, and competition on climate-driven range shifts in trees. PLoS One 1011: e0142369.

Naugžemys D, Lambertini C, Patamsytė J, Butkuvienė J, Khasdan V, Žvingila D. 2021. Genetic diversity patterns in Phragmites australis populations in straightened and in natural river sites in Lithuania. Hydrobiologia 848: 3317–3330.

Nei M. 1987. Molecular evolutionary genetics. Columbia university press.

Noguerales, V, García-Navas, V, Cordero, PJ, Ortego, J. 2016. The role of environment and core-margin effects on rangewide phenotypic variation in a montane grasshopper. Journal of Evolutionary Biology, 29(11), 2129–2142.

Nybom H. 2004. Comparison of different nuclear DNA markers for estimating intraspecific genetic diversity in plants. Molecular ecology 135: 1143–1155.

Orsini L, Vanoverbeke J, Swillen I, Mergeay J, De Meester L. 2013. Drivers of population genetic differentiation in the wild: isolation by dispersal limitation, isolation by adaptation and isolation by colonization. Molecular ecology 2224: 5983–5999.

Padgett DJ. 2007. A MONOGRAPH OF NUPHAR (NYMPHAEACEAE). Rhodora 109: 1–95.

Palmer MA, Hakenkamp CC, Nelson-Baker K. 1997. Ecological heterogeneity in streams: why variance matters. Journal of the North American Benthological Society 161: 189–202.

Pierce S, Brusa G, Sartori M, Cerabolini BEL. 2012. Combined use of leaf size and economics traits allows direct comparison of hydrophyte and terrestrial herbaceous adaptive strategies. Annals of Botany 1095: 1047–1053.

Price TD, Qvarnström A, Irwin DE. 2003. The role of phenotypic plasticity in driving genetic evolution. Proceedings of the Royal Society of London Series B: Biological Sciences 270: 1433–1440.

Puijalon S, Bouma TJ, Douady CJ, et al. 2011. Plant resistance to mechanical stress: evidence of an avoidance–tolerance trade-off. New Phytologist 1914: 1141–1149.

Qiu T, Jiang LL, Yang YF. 2016. Genetic and epigenetic diversity and structure of Phragmites australis from local habitats of the Songnen Prairie using amplified fragment length polymorphism markers. Genet Mol Res 15.

Rasband WS. 2018. WS 1997–2018 ImageJ US National Institutes of Health, Bethesda, Maryland, USA.

Reale L, Gigante D, Landucci F, Venanzoni R, Ferranti F. 2011. Correlation between sexual reproduction in Phragmites australis and die-back syndrome. The International Journal of Plant Reproductive Biology 32: 133–140.

Ren L, Guo X, Liu S, et al. 2020. Intraspecific variation in Phragmites australis: Clinal adaption of functional traits and phenotypic plasticity vary with latitude of origin. Journal of Ecology 1086: 2531–2543.

Reynolds J, Weir BS, Cockerham CC. 1983. Estimation of the coancestry coefficient: basis for a short-term genetic distance. Genetics 1053: 767–779.

Richards CL, Schrey AW, Pigliucci M. 2012. Invasion of diverse habitats by few Japanese knotweed genotypes is correlated with epigenetic differentiation. Ecology letters 159: 1016–1025.

Rogers AR, Harpending H. 1992. Population-growth makes waves in the distribution of pairwise genetic-differences. Molecular Biology and Evolution 9: 552–569.

Sawilowsky SS. 2009. New effect size rules of thumb. Journal of modern applied statistical methods 8: 26.

Schneider S, Roessli D, Excoffier L. 2000. Arlequin: a software for population genetics data analysis. User manual ver 2: 2496–2497.

Schoelynck J, Bal K, Verschoren V, et al. 2014. Different morphology of Nuphar lutea in two contrasting aquatic environments and its effect on ecosystem engineering. Earth surface processes and landforms 39: 2100–2108.

Seymour M, Räsänen K, Kristjánsson BK. 2019. Drift versus selection as drivers of phenotypic divergence at small spatial scales: The case of Belgjarskógur threespine stickleback. Ecology and Evolution 9: 8133–8145.

Slatkin M. 1995. A measure of population subdivision based on microsatellite allele frequencies. Genetics 139: 457–462.

Spafford L, Le Maire G, MacDougall A, De Boissieu F, Féret JB. 2021. Spectral subdomains and prior estimation of leaf structure improves PROSPECT inversion on reflectance or transmittance alone. Remote Sensing of Environment 252: 112176.

Spitze K. 1993. Population structure in Daphnia obtusa: quantitative genetic and allozymic variation. Genetics 135: 367–374.

Takeda S, Kurihara Y. 1988. The effects of the reed, Phragmites australis Trin, on substratum grain-size distribution in a salt marsh. Journal of the Oceanographical Society of Japan 443: 103–112.

Team RC. 2019. R: a language and environment for statistical computing, version 3.0. 2. Vienna, Austria: R Foundation for Statistical Computing, 2013.

Thomaz SM. 2021. Ecosystem services provided by freshwater macrophytes. Hydrobiologia 1–21.

Villa P, Bolpagni R, Pinardi M, Tóth VR. 2021. Leaf reflectance can surrogate foliar economics better than physiological traits across macrophyte species. Plant Methods 171: 1–16.

Villa P, Pinardi M, Tóth VR, Hunter PD, Bolpagni R. 2017. Remote sensing of macrophyte morphological traits: implications for the management of shallow lakes. Journal of Limnology 76(1): 109–126.

Vretare V, Weisner SE, Strand JA, Granéli W. 2001. Phenotypic plasticity in Phragmites australis as a functional response to water depth. Aquatic Botany 69, 127–145.

Vymazal J. 2011. Plants used in constructed wetlands with horizontal subsurface flow: a review. Hydrobiologia 674: 133–156.

Vyšniauskienė R, Rančelienė V, Naugžemys D, et al. 2020. Genetic diversity of Nuphar lutea in Lithuanian river populations. Aquatic Botany 161: 103173.

Waddington CH. 1953. Genetic assimilation of an acquired character. Evolution 7: 118–126.

Waddington CH. 1952. Selection of the genetic basis for an acquired character. Nature 169: 625–626.

Wang R, Gamon JA. 2019. Remote sensing of terrestrial plant biodiversity. Remote Sensing of Environment 231: 111218.

Wellburn AR. 1994. The spectral determination of chlorophylls a and b, as well as total carotenoids, using various solvents with spectrophotometers of different resolution. Journal of plant physiology 1443: 307–313.

West-Eberhard MJ. 2003. Developmental plasticity and evolution. Oxford University Press.

Whitlock MC. 2008. Evolutionary inference from QST. Molecular ecology 178: 1885–1896.

Yang AH, Wei N, Fritsch PW, Yao XH. 2016. AFLP genome scanning reveals divergent selection in natural populations of Liriodendron chinense Magnoliaceae along a latitudinal transect. Frontiers in plant science 7: 698.

